# Epigenetically Modified DNAzyme Walkers for Simultaneously Imaging of MGMT and FTO Activity in Living Cancer Cells

**DOI:** 10.1101/2025.06.15.659742

**Authors:** Yewei Li, Liang Fuling, Mingda Li, Yanfang Gong, Die Long, Huanhuan Sun, Min Hou, Mingyan Jiang

## Abstract

Investigations into epigenetic modifications have underscored the pivotal roles demethylases play in modulating the dynamics of RNA and DNA. Abnormal demethylase expression is closely related to the occurrence and development of many diseases. Unraveling the individual activity of different demethylases is essential to study the underlying mechanisms. In this paper, we report the development of demethylase-responsive DNAzyme-powered walkers designed for the simultaneous imaging of two important demethylases: O6-methylguanine-DNA methyltransferase (MGMT) and fat mass and obesity-associated protein (FTO). Epigenetically inactivated DNAyzme-based DNA walker-functionalized gold nanoparticles were employed to orthogonally visualize MGMT and FTO activities in various live cancer cell lines with high sensitivity. This method offers a direct way for the comprehensive evaluation of multiple demethylase activities, and the understanding of epigenetic regulation.

## Introduction

Recent investigations into the realm of epigenetics have elucidated that modifications such as methylation, acetylation, and the incorporation of pseudouridine in DNA and RNA serve pivotal roles in a multitude of cellular mechanisms^1,2^. Among these, the processes of methylation and demethylation, governed by their respective “writer” and “eraser” enzymes, represent the most prevalent forms of epigenetic regulation^3, 4^. Of particular interest are DNA demethylases, a category of enzymes tasked with the removal of methyl groups from modified nucleic acids^5^. Dysregulation in the activity of these demethylases has been tightly linked with aberrant gene expression patterns, decreased responsiveness to therapeutic interventions, and enhanced progression of neoplastic diseases^6, 7^. For example, the upregulation of the initially identified demethylase, FTO, has been implicated as a significant participant in oncogenesis through its role in m6A demethylation, underscoring the crucial implications of epigenetic modifications in both health and disease^8-10^.

Therefore, methods for profiling demethylase activity in living cells would greatly enhance understanding of epigenetic regulatory mechanisms significantly^11-14^. Recently, several groups have reported the development of DNAzyme-based biosensors for the detection of demethylase activity. For example, Wang et al. have screened the 8-17 DNAzyme with N6-methyladenine (m6A) substitution, which could be selectively reactivated by FTO, thereby enabling the intracellular imaging of FTO activity and its regulatory effects on gene expression^15^. Similar approaches were further demonstrated by a ‘repaired and activated’ DNAzyme that responds to MGMT activity^16^. However, these approaches primarily depend upon the reactivation of DNAzymes to hybridize with freely diffusing substrates to induce cleavage and subsequently fluorescence recovery, providing poor signal-to-noise ratios (SNRs) for the imaging of demethylase activity. To address this, Huang et al. reported the development of a cascade amplification, integrating DNAzyme-CRISPR/cas12a, aimed at enhancing the detection of MGMT activity^17^. However, it is crucial to acknowledge that this strategy is not applicable for the profiling of live cell demethylase activity, underscoring the necessity for further methodological developments in this domain.

To establish a highly sensitive platform for imaging intracellular demethylase activity, we propose using a DNAzyme-powered DNA walker, which can efficiently amplify localized signals. The modular integration of DNAzymes within DNA walkers enables the activation of DNA walkers, enhancing their responsiveness^18, 19^. DNAzyme-based walkers have been effectively demonstrated as self-powered, in situ signal amplifiers on nanoscale surface^20, 21^. For instance, Le’s group developed a DNAzyme walker for detecting apurinic/apyrimidinic endodeoxyribonuclease 1 (APE1) in live cells, achieving a detection limit (LOD) of 160 fM22. Additionally, Ju’s group introduced a DNAzyme walker-based imaging method for cooperative visualization of mutant p53 and telomerase^23^.

## Results

**Figure 1A** presents the schematic of our demethylase-activated DNAzyme approach, utilizing the 8-17E DNAzyme caged with N6-methyladenine (17E-Me) and O6-methylguanine (O6MeG) as a model system. Initially, we evaluated the performance of 17E-Me and O6MeG EMOzymes. As shown in **Figure 1B&C**, these demethylase-activated DNAzymes specifically induced substrate cleavage without cross-reactivity between MGMT and FTO. Metal ion dependence testing further revealed that these EMOzymes did not interfere with each other **(Figures 1D&E)**. These results confirm the feasibility of using 17E-Me and O6MeG EMOzymes for orthogonal detection of MGMT and FTO.

**Figure 1.**
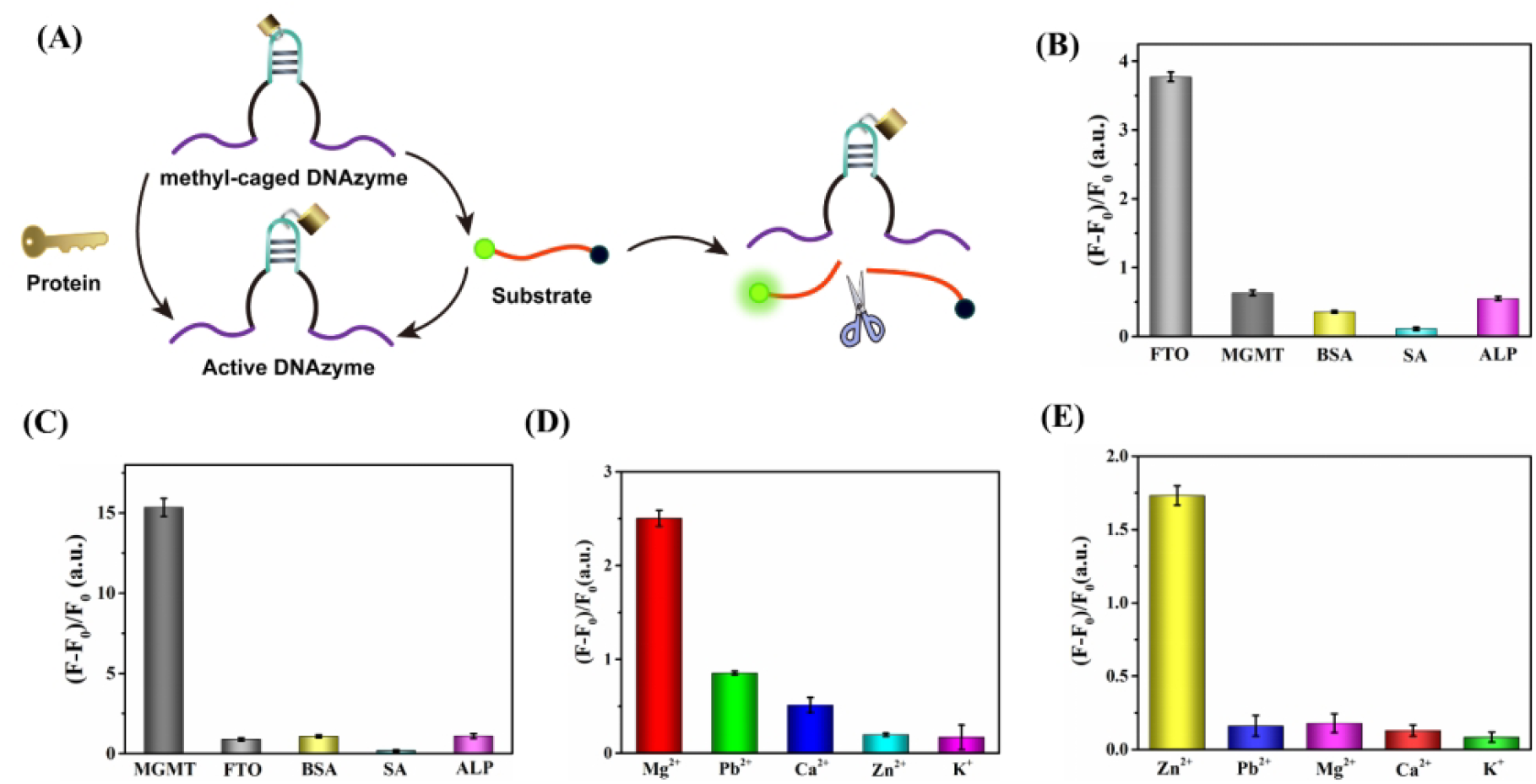
Specific activation of the 17E-Me and O6MeG EMOzyme by FTO and MGMT. **(A)** Diagram illustrating demethylase-activated EMOzymes through efficient protein-mediated demethylation of DNA; **(B)** Fluorescence enhancement of the 17E-Me EMOzyme incubated with various proteins; **(C)** Fluorescence response of the O6MeG EMOzyme treated with various proteins; **(D)** Fluorescence response of the O6MeG EMOzyme treated with different metal ions; **(E)** Fluorescence response of the 17E-Me DNAzyme treated with different metal ions.、

We next designed the 17E-Me and O6MeG EMOWAs and functionalized them onto gold nanoparticles (AuNPs) to create dual-EMOWA@AuNP probes for the sensitive detection of MGMT and FTO **(Figure 2A)**. The size, absorption spectrum, and zeta potential of the AuNP and EMOWA@AuNP probes were characterized using TEM, UV-VIS spectroscopy, DLS, and a zetasizer **(Figures 2B-E)**. The AuNPs exhibited uniform dispersion, and successful EMOWA modification was indicated by an absorption peak at 280 nm, increased DLS size. We then quantitatively measured demethylase-dependent fluorescence recovery from EMOWA@AuNP **(Figure 2F&H)**. Emission intensities of FITC at 525 nm and 580 nm were plotted against demethylase concentrations, showing good linearity, which suggests that the EMOWA@AuNP response is proportional to demethylase concentration. The linear detection ranges for 17E-Me EMOWA@AuNP and O6MeG EMOWA@AuNP were 0-200 nM and 0-250 nM, respectively **(Figures 2G&I)**. The calculated limits of detection (LOD) were 3.02 nM for MGMT and 0.93 nM for FTO, yielding fluorescence signals distinguishable from the background and demonstrating high sensitivity.

**Figure 2.**
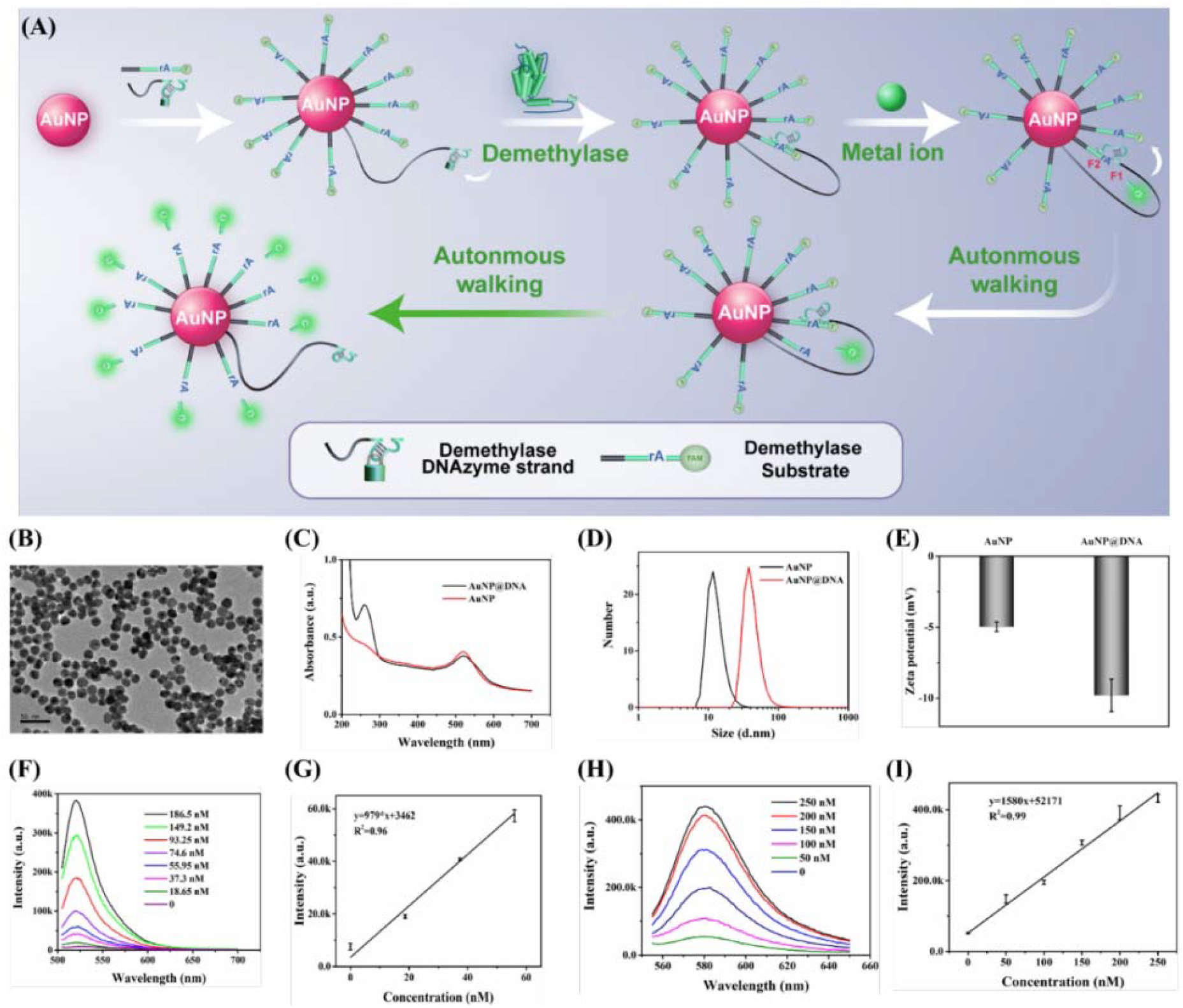
**(A)** Working mechanism of AuNP-loaded DNAzyme probes for in vitro demethylase detection. **(B)** Transmission electron microscopy (TEM) images of EMOWA@AuNP. **(C)** UV-vis absorption spectra, **(D)** Dynamic light scattering (DLS) characterization, and **(E)** Zeta potential of AuNPs and EMOWA@AuNP. **(F)** The fluorescence response of the O6MeG EMOWA@AuNP walking motor was analyzed with various concentrations of the MGMT protein. **(G)** Calibration curves for the O6MeG-mediated MGMT assay were constructed by plotting fluorescence intensity against MGMT concentration. **(H)** The fluorescence response of 17E-Me EMOWA@AuNP was analyzed with various concentrations of FTO protein. **(I)** Calibration curves for the 17E-Me-mediated FTO assay were constructed by plotting fluorescence intensity against FTO concentration.

The DNAzyme motor was applied to operate within T98G human glial cells to evaluate its intracellular functionality. MGMT and FTO, the target proteins, are expressed at low concentrations in this cell line, presenting substantial challenges for imaging using conventional methods. To facilitate cellular uptake, the DNAzyme motor and its track were functionalized on AuNPs, as DNA-functionalized AuNPs have been shown in previous studies to enter cells effectively without requiring transfection reagents. Additionally, two negative control probes were designed, incorporating mismatches in the DNAzyme’s catalytic and substrate-binding domains (designated as NC-Dz and NC-S1). These control probes lacked catalytic activity, ensuring that any observed fluorescence originated exclusively from catalytic cleavage by the active DNAzyme, rather than from nonspecific target binding. Both EMOWA@AuNP and control nanoprobes (NC-Dz@AuNP and NC-S1@AuNP) were sequentially incubated with T98G cells and analyzed using confocal laser scanning microscopy (CLSM). As illustrated in **Figures 3A&C**, no fluorescence was detected in cells treated with control nanoprobes, confirming the stability of the substrate strand on AuNPs in the absence of an active DNAzyme motor. Fluorescence was observed only when Mg^2+^ and O^6^MeG EMOWAs@AuNP nanoprobes were both present, yielding green fluorescence in T98G cells, and when Zn^2+^ and 17E-Me EMOWAs@AuNP nanoprobes coexisted, producing red fluorescence, thereby validating the successful operation of the DNAzyme motor within living cells. Consistent results were obtained using flow cytometry (**Figures 3E&G**), demonstrating the effectiveness of the EMOzyme-powered DNA walker for imaging MGMT and FTO fluorescence in live cells.

**Figure 3.**
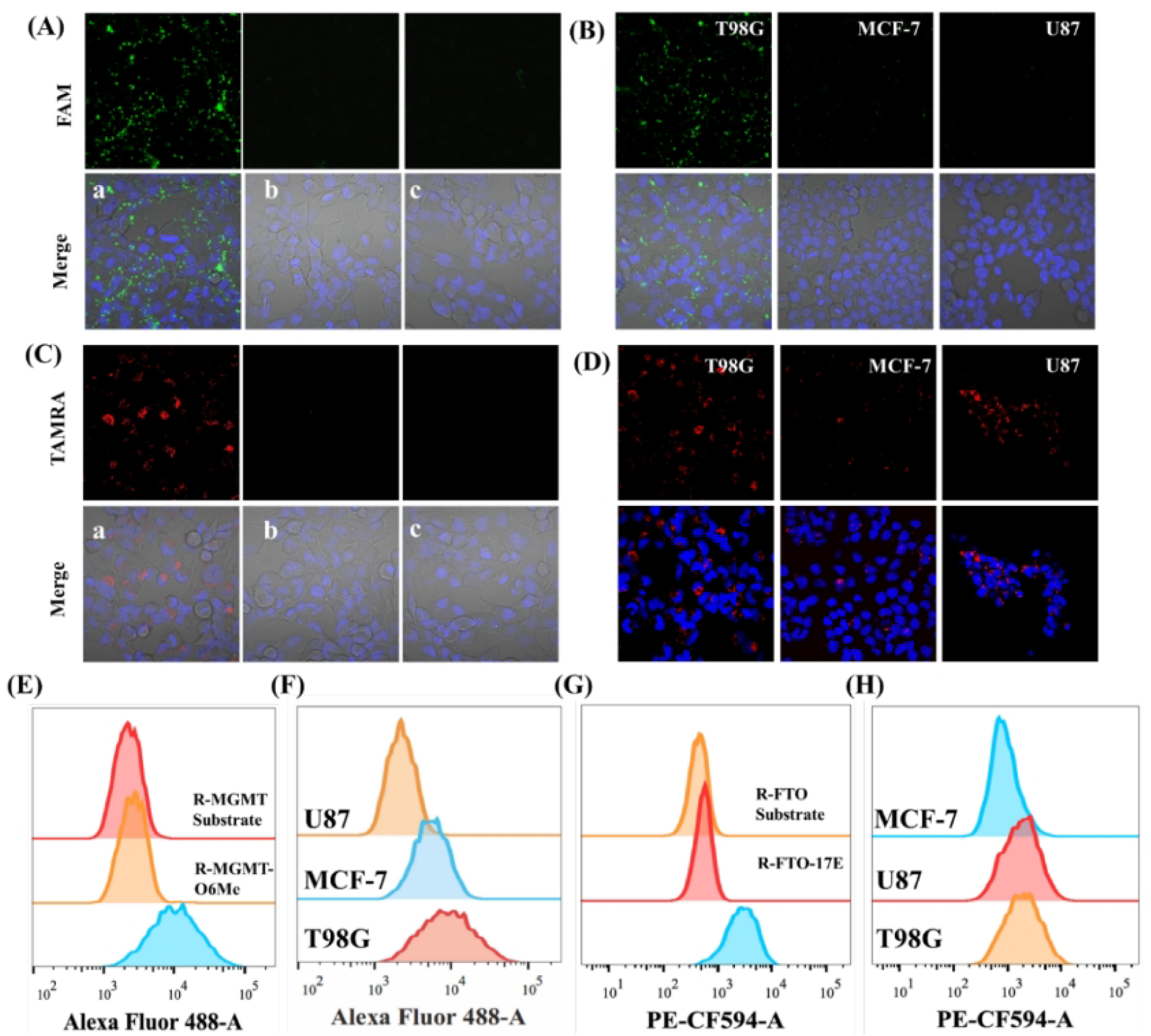
EMOWA@AuNP-based fluorescence analysis of MGMT and FTO in live cells. **(A)** Fluorescence imaging of endogenous MGMT in live cells was conducted as follows: T98G cells treated with **(a)** O6MeG DNAzyme@S1@AuNP, **(b)** NC-Dz@S1@AuNP, and **(c)** O6MeG DNAzyme@NC-S1@AuNP. **(B)** Confocal fluorescence imaging of intracellular MGMT activity was performed in T98G, MCF-7, and U87 cells (scale bar: 20 μm). **(C)** Fluorescence imaging of endogenous FTO in live cells was conducted as follows: **(a)** T98G cells treated with 17E-Me DNAzyme@S2@AuNP, **(b)** T98G cells treated with NC-Dz@S2@AuNP, and (c) T98G cells treated with 17E-Me DNAzyme@NC-S2@AuNP. **(D)** Fluorescence imaging of intracellular FTO activity was performed in T98G, MCF-7, and U87 cells (scale bar: 20 μm). **(E)** Flow cytometric analysis of endogenous MGMT in T98G cells was conducted. **(F)** Flow cytometric analysis of endogenous MGMT in various cell types was conducted. **(G)** Flow cytometric analysis of endogenous FTO in T98G cells was performed. (H) Flow cytometric analysis of endogenous FTO in various cell types was conducted.

To further examine the specificity and general applicability of the nanoprobes, EMOWAs@AuNP nanoprobes were tested on three cancer cell lines: T98G (human glioblastoma), MCF-7 (breast cancer), and U87 (human glioma). Each cell type was incubated with O^6^MeG EMOWAs@AuNP and 17E-Me EMOWAs@AuNP for 8 hours and subsequently imaged using CLSM. As shown in **Figure 3B**, T98G cells displayed strong fluorescence for MGMT, indicating high expression levels of this target, whereas MCF-7 and U87 cells showed only weak fluorescence, suggesting low MGMT expression. **Figure 3D** shows that T98G and U87 cells displayed strong fluorescence for FTO, indicating high expression of this protein, while MCF-7 cells exhibited weaker fluorescence, suggesting lower FTO expression.

## Discussion

In conclusion, we developed EMOzyme-powered DNA walker-functionalized gold nanoparticles (AuNPs) to simultaneously visualize the active expression of MGMT and FTO in living cells. This nanoprobe specifically activates upon recognizing MGMT and FTO, with the DNAzyme walking strands autonomously cleaving the corresponding substrates arranged on the nanoprobe surface in the presence of metal ions, thereby producing numerous fluorescent fragments. We successfully applied the demethylase-activated DNAzyme strategy for real-time multiplex visualization of MGMT and FTO in living cells and a tumor-bearing mouse model. This study highlights the feasibility of utilizing EMOWA to report multiple demethylase activities, providing a valuable method for visualizing demethylase activity in epigenetic research.

## Methods

### Cell Culture

Cells (MCF-7, T98G, U-87) were thawed, centrifuged, and resuspended in DMEM (Sigma-Aldrich) supplemented with 10% FBS (Sigma-Aldrich) and antibiotics. Cultures were maintained at 37°C/5% CO□. At 80% confluency, cells were passaged using trypsin-EDTA and subcultured.

### Live Cell Imaging

Cells (85% confluency) in confocal dishes were incubated for 5 h in Opti-MEM (Sigma-Aldrich) containing DNAzyme motors and 10 mM Mg^2^□. After washing, cells were cultured in complete medium for 12 h prior to imaging (Olympus confocal system).

### Polyacrylamide Gel Electrophoresis

Native PAGE gels were prepared using acrylamide/bis-acrylamide and TBE buffer. Samples were electrophoresed at 120 V for 60 min. Gels were stained with GelRed (Tsingke Biotech) and imaged (gel documentation system).

### Demethylase Activity Assay

MGMT System: O□-methylated DNAzyme (500 nM) and MGMT (250 nM) in Buffer B were incubated overnight at 37°C. Substrate S1 (500 nM) was added for 5 h before fluorescence measurement. FTO System: m□A-modified DNAzyme (500 nM) and FTO (250 nM) in Buffer A were similarly incubated, followed by S2 addition. Specificity: FAM/TAMRA-labeled probes (500 nM) were incubated with target/non-target proteins, with fluorescence monitored.

### Cytotoxicity Assay (CCK-8)

T98G cells (5×10^3^/well) were treated with EMOWA@AuNP/metal ions. After incubation, WST-8 reagent (Abbkine) was added and absorbance measured (Molecular Devices microplate reader, n=3).

### Cellular Analysis

Fluorescence Imaging: Cells treated with Zn^2^□/Mg^2^□ and EMOWA@AuNP probes were nuclear-stained (Hoechst 33342) and imaged. Western Blot: Lysates were quantified (BCA kit, Tsingke Biotech), separated by SDS-PAGE, transferred to PVDF membranes, and probed with primary/secondary antibodies. Detection used ECL reagent (Abbkine).

